# DESIGNING OF CUSTOM BARCODES FOR SEQUENCING ON THE MGI PLATFORM

**DOI:** 10.1101/2022.09.07.506907

**Authors:** Irina Bulusheva, Anna Shmitko, Julia Vasiliadis, Oleg Suchalko, Denis Syrko, Vera Belova, Dmitriy Korostin

**Affiliations:** Center for Precision Genome Editing and Genetic Technologies for Biomedicine Pirogov Russian National Research Medical University, Ostrovityanova str. 1, Moscow, Russian Federation, 117997

**Keywords:** DNBSEQ, MGI, BGI, NGS, custom barcodes, custom indexes, custom adapters

## Abstract

The recently launched MGI platform (DNBSEQ-G50, -G400 and -T7 sequencers) has been gaining popularity for next-generation sequencing. However, the barcode set for library preparation provided by the manufacturer has certain limitations on the number of samples that can be sequenced at a time and the compatibility of barcodes from different or incomplete sets as well as a ratio between the samples.

In this paper, we present a protocol for designing custom barcodes expanding the pre-existing barcode set and demonstrate its performance on the MGI Tech Co. Ltd. (China) machines. We developed a universal “quad method” which allows for simultaneous sequencing of any number of samples up to 252 per lane which is multiple of 4 or 4n+2. Here, we describe this method, its analysis, verification, and integration into the sequencing as well as its validation for sequencing using the DNBSEQ G-400 machine.

## Introduction

There are quite a few of custom solutions developed for various applications employing the platform for mass parallel sequencing Illumina ^1,2,3,4^. An alternative to Illumina platform is provided by the MGI Tech Co. Ltd. company (a subsidiary of the BGI Group) which produces the range of sequencers based on the DNA nanoball technology and cPAS sequencing. It allows for sequencing in the single-end or paired-end mode using single or dual barcode conditions. The technology presupposes barcoding of samples during ligation of the adapters containing barcode sequences. DNA library barcoding is necessary for labeling sequences from different biological samples and read identification during the transformation of temporary sequencing files into the commonly used fastq format. The length of MGI barcodes is 10 bp.

By default, the standard kits for library preparation and sequencing with the mid-throughput sequencer DNBSEQ G-400 are designed for a single-index sequencing, whereas the dual barcode mode is optional and requires purchasing additional kits. Currently, MGI provides the kits that include 96 barcode adapters https://en.mgi-tech.com/products/reagents_info/5/ and 128 sequences for synthesis (https://en.mgi-tech.com/Download/download_file/id/71).

The G-400 system is sensitive to the nucleotide balance at each cycle of barcode sequencing as the quality drastically drops if the same position in the barcode sequences from the same lane is occupied by the same nucleotide. This explains why the barcode set from the same lane should meet the criteria for combining their sequences and enable generating compatible sets. The set of 96 adapters provided by MGI allows for forming 11 sets. In practice, however, it is often necessary to combine the samples containing barcodes from different sets, change the number of the samples loaded on a lane and to vary their ratio. Therefore, the manufacturer imposes limitations on the users of this platform providing a small number of barcodes and sets, which thus prevents uncovering its true potential for sequencing. This may prove critical when selecting a sequencing platform.

Earlier, we developed the software allowing for designing the optimal combination of provided barcodes at various ratios and sample numbers^5^. The updated software including custom barcodes is available at the link (https://github.com/genomecenter/BC-store).

In this work, we developed an algorithm that can be used for generating the necessary number of barcode sequences required for a study. Using this algorithm, we designed 252 barcodes forming 63 balanced sets each consisting of 4 barcodes and allowing any set to be combined with the others.

## Method concept and barcode design

### The ‘quad method’ algorithm

The sequencer has limits in terms of intensity of the registered signal from the fluorophores corresponding to the nucleotides. If the same position of barcodes contains the same nucleotide, read quality significantly drops leading to the errors in barcode identification and further assigning reads to the samples ^5^. Therefore, we had to design barcodes to generate the most balanced combinations. The algorithm of sequence design is based on the ‘quad method’ that involves adding 3 barcodes obtained by the consecutive substitutions of bases to each barcode from the MGI set (Fig.1A, 1B).

**Figure 1.**
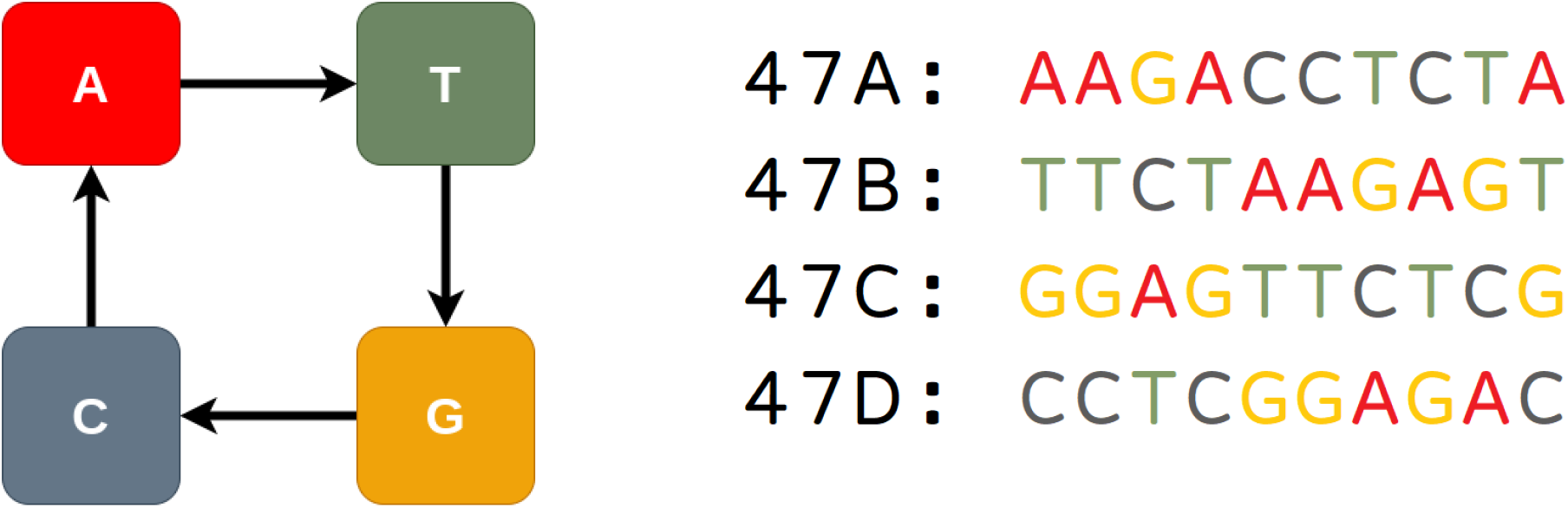
A - the concept of the quad method. Each MGI barcode serves as a root for the quad, custom barcodes being generated by sequential changes at each position of the original barcode: A->T, T->G, G->C, C->A. B - the example of a barcode quad. 47A is the original MGI barcode. 47B, 47C, 47D are the custom barcodes generated using the quad method.

Following this method, each of 96 barcodes can serve as a root barcode for its quad resulting in generating 86*4=384 unique barcodes.

As the percentage of each base at each position is 25%, the resulting combination is perfectly balanced and guarantee highest quality of sequencing.

### Verification of compliance with the criteria

#### Verifying the compatibility of barcodes based on a mismatch number

At the next step, all quads were checked for the compatibility based on the mismatch number. Each sample labeled by a barcode had to be uniquely identified, so the barcode sequences of certain length should not overlap with the others. We selected a threshold of 4 mismatches as all 96 10 bp barcodes provided by the manufacturer differ by more than 4 bases. We constructed a graph of incompatible quads (S1 Fig.) and using an adjacency matrix (S2 Fig.), we selected 63 quads (252 barcodes) compatible between each other based on the number of permitted mismatches (Fig.2). The sequences of all 252 barcodes are listed in S1 Table.

**Figure 2.**
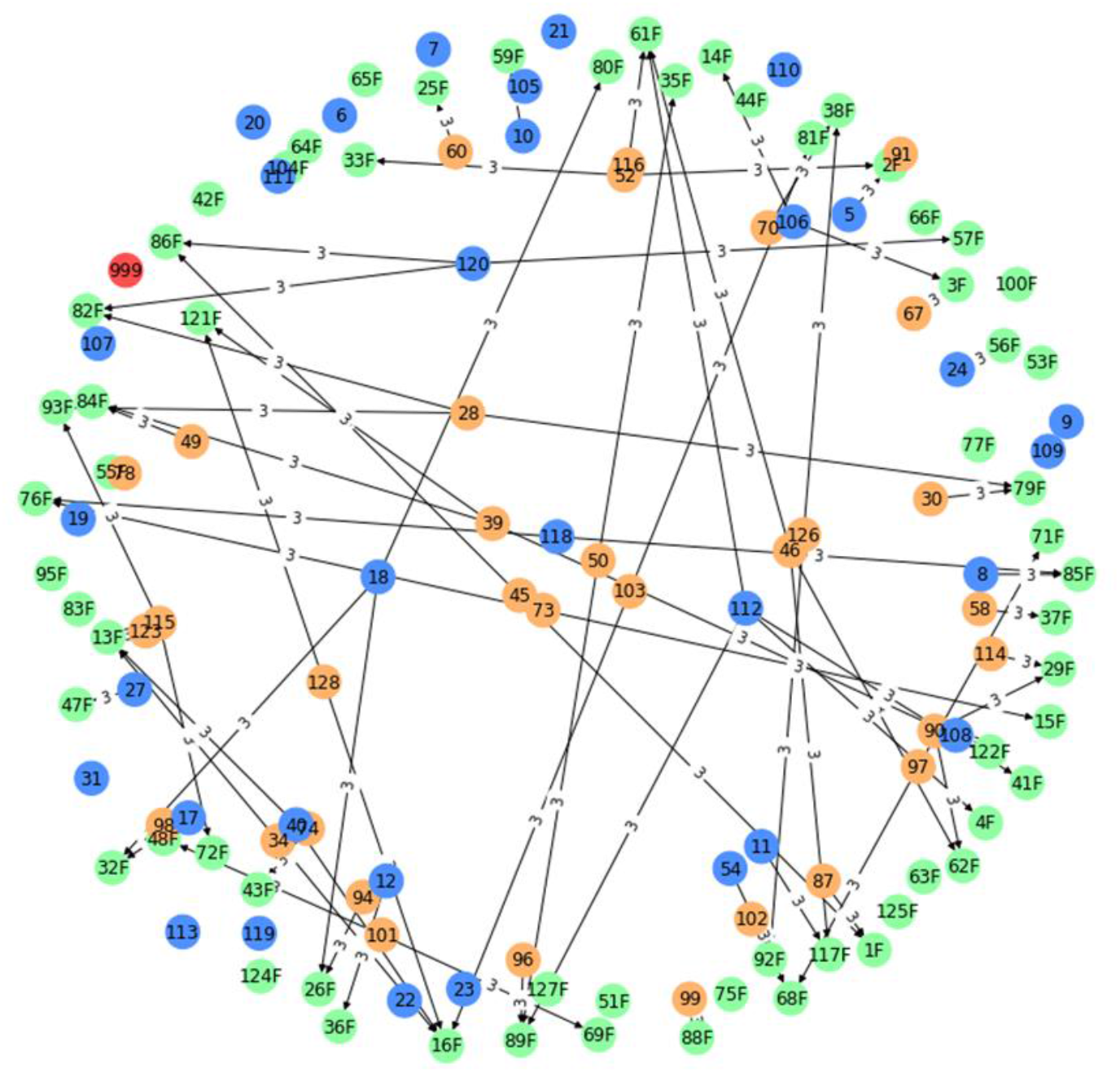
The incompatibility graph showing 63 quads and MGI barcodes not included in the quads. All quads that passed filtering based on the mismatch number are shown in green, the barcodes from the set of 96 MGI barcodes not included in the quads are shown in orange, the MGI barcodes from the set of 128 MGI barcodes are shown in blue, the 999 verification MGI barcode is shown in red. The line connects the incompatible barcodes and quads; the number above the line indicates the lowest number of mismatches between them.

### Validation of the compatibility based on the balance

As each quad is perfectly balanced, any number of quads can be combined between each other. The ratio between the quads in a pool can vary; however, ratios between barcodes in each quad should be equal.

Furthermore, we checked if it was possible to generate the pools containing 4n+2 barcodes, where n is the number of quads. We checked the compatibility using the combination of 10 barcodes by the BC-Store software (Fig.3). Nucleotide fraction of each nucleotide at any position in a pool of 10 barcodes has a highest and lowest deviations equal to 0.2 and 0.3, respectively, and meets the conditions of strong criteria indicating a balanced combination. This is still valid when any of the two barcodes from the same quad are added to n quads at a ratio equal or lower than in quads.

**Figure 3.**
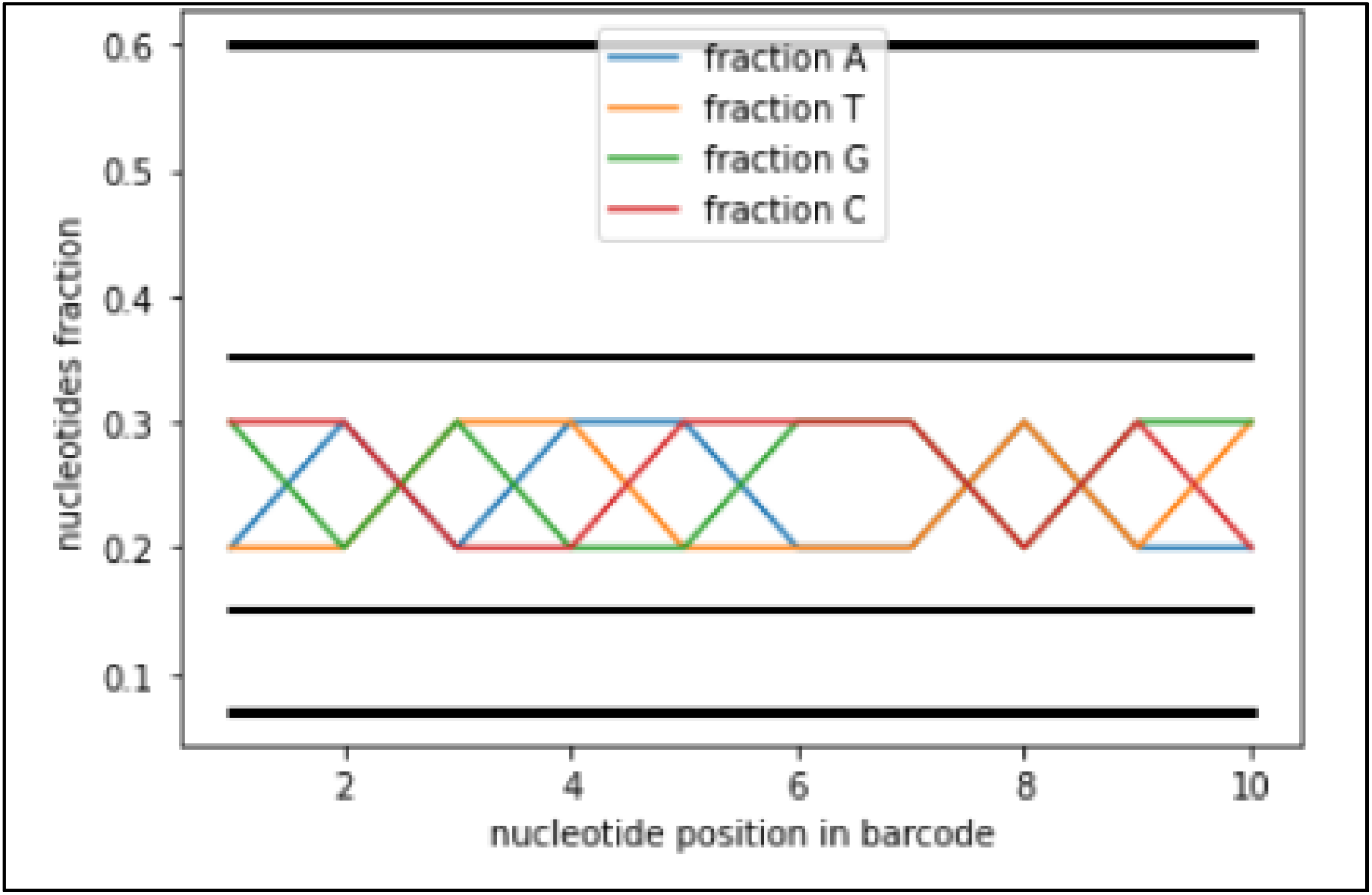
Nucleotides balance in 4n+2 barcodes pool. Colored lines represent the nucleotide fractions. Black strong lines represent the boundaries of the lite criterion, fine lines represent the boundaries of the strong criterion of barcode compatibility ^5^.

### Validating the uniqueness

We checked if the sequences of designed barcodes are present among the original MGI barcodes. This is necessary for generating the file containing barcodes for automatic demultiplexing. For this purpose, we created a Venn diagram showing the sets of custom and original MGI barcodes. We obtained 63 overlaps, where all 63 barcodes were original MGI barcodes, while other 189 sequences were unique sequences not coinciding with the MGI barcodes from different kits.

## Results

### Adapter synthesis

According to the manufacturer’s instructions, designing an individual adapter requires annealing two oligonucleotides. One of them (top oligonucleotide) contains the barcode sequence and a phosphate at the 5’-end (Ad153_5T_1-index # (1∼128) according to the manufacturer), the sequence of bottom oligonucleotide is partially complementary to the top oligonucleotide (https://en.mgitech.cn/Download/download_file/id/71). These oligonucleotides form “bubble” adapter during annealing in PCR tube suitable to produce adapter-flanked DNA library via TA-ligation reaction with T4 DNA ligase.

The sequences of oligonucleotides containing the barcodes 1A-1D are shown in Table 1, all sequences containing 252 barcodes are listed in S1 Table.

**Table 1.**
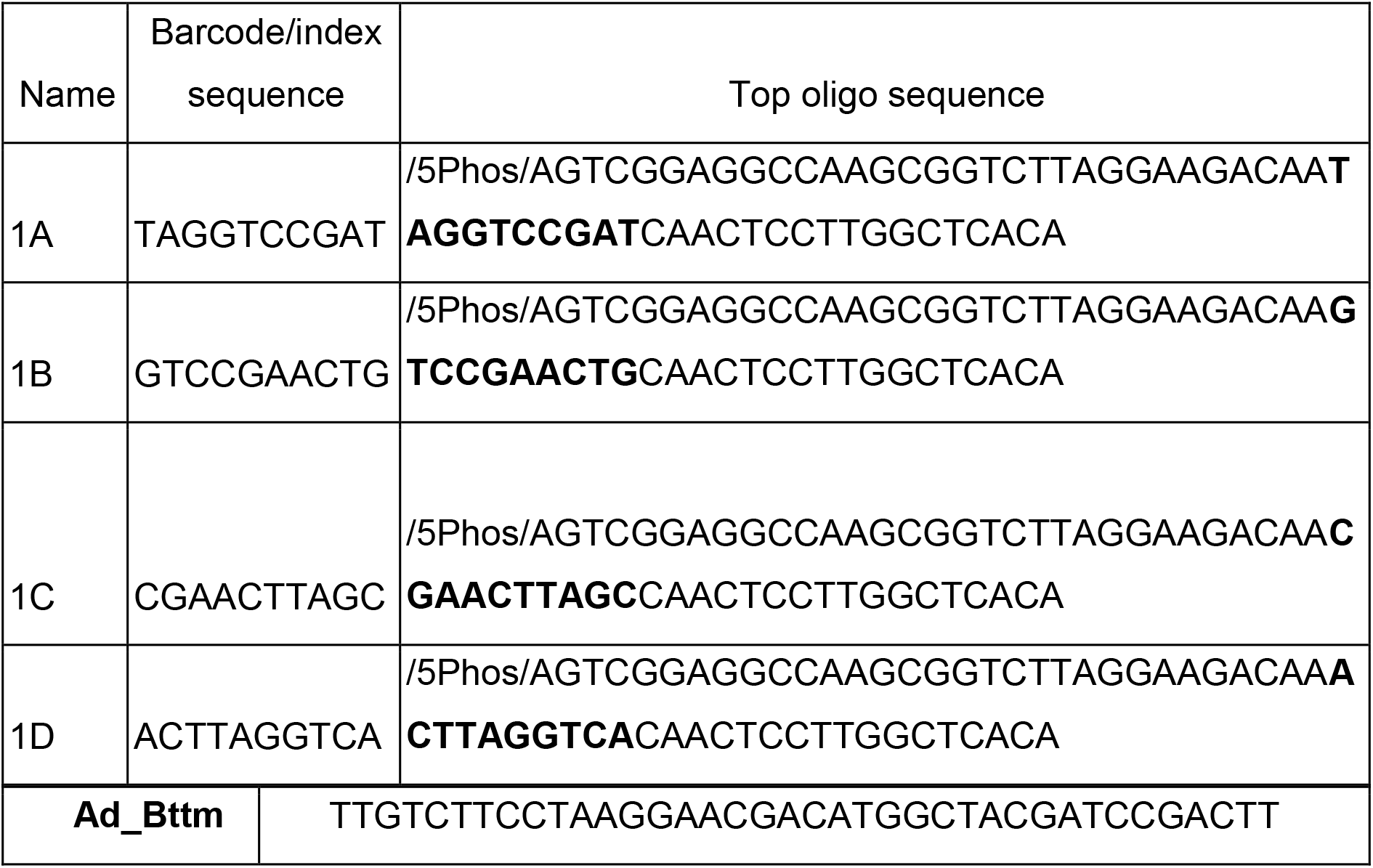
The list of barcode sequences and full sequences of top and bottom oligonucleotides for adapter preparation. Ad_Bttm -bottom oligo.

Adapters were prepared as follows:

1 ul of 5M NaCl, 10 ul of 200 uM top oligo, and 10 ul of 200 uM bottom oligo were added to 70 ul of the LowTE buffer. The mix was heated at 95°C for 2 min and gradually chilled to 17°C by 0.5°C every 30 s.

### The algorithm of uploading new barcodes to a sequencer

For automatic demultiplexing of the sequenced libraries and following the recommendations of MGI, we created a .csv file (S2 Table) with barcode sequences including new custom barcodes, the original MGI barcodes from the 96 and 128 kits and 999 barcodes from MGI. The MGI barcodes included into the quads had a nA structure, where n is an adapter number in the original MGI kit, while custom barcodes had nB, nC, nD structure according to the order of quad formation. The format of the original MGI barcodes not included in quads remained as before. The barcode numbers were separated from the barcode sequences with commas without spaces.

### Validation of libraries with the custom barcodes on DNBSEQ G-400

To validate the designed barcodes, we prepared the libraries with the synthesized custom adapters (Evrogen). The libraries, prepared following the standard MGI protocol, were pooled and enriched using the SureSelect Human All Exon v7 kit and then sequenced in the PE100 mode using the DNBSEQ G-400 machine. Fastq demultiplexing was performed by the software built in G-400 provided by MGI basecalllite based on the uploaded file containing the barcode sequencing data. By default, the algorithm considers a read “undecoded” if there are two or more mismatches in a 10 bp barcode sequence. Therefore, the fraction of undecoded reads can be used as a quality metrics of the performance of DIY barcode adapter. We compared the fraction of undecoded reads in complete data from each lane (in Gb) with custom barcodes (4 lanes) with the data from 10 previous runs with MGI barcodes. On average (mean±SD), the fractions of undecoded reads per lane were 1.08± 0.0019% and 1.72± 0.002% for the MGI adapters and custom adapters, respectively (Fig.5). We assume that a higher value of undecoded reads may be related to the lower quality of the synthesized oligonucleotides compared to MGI. Although the difference in the ratio of undecoded reads is significant, we consider the absolute value relative to the total data output from a single track to be negligible.

**Figure 4.**
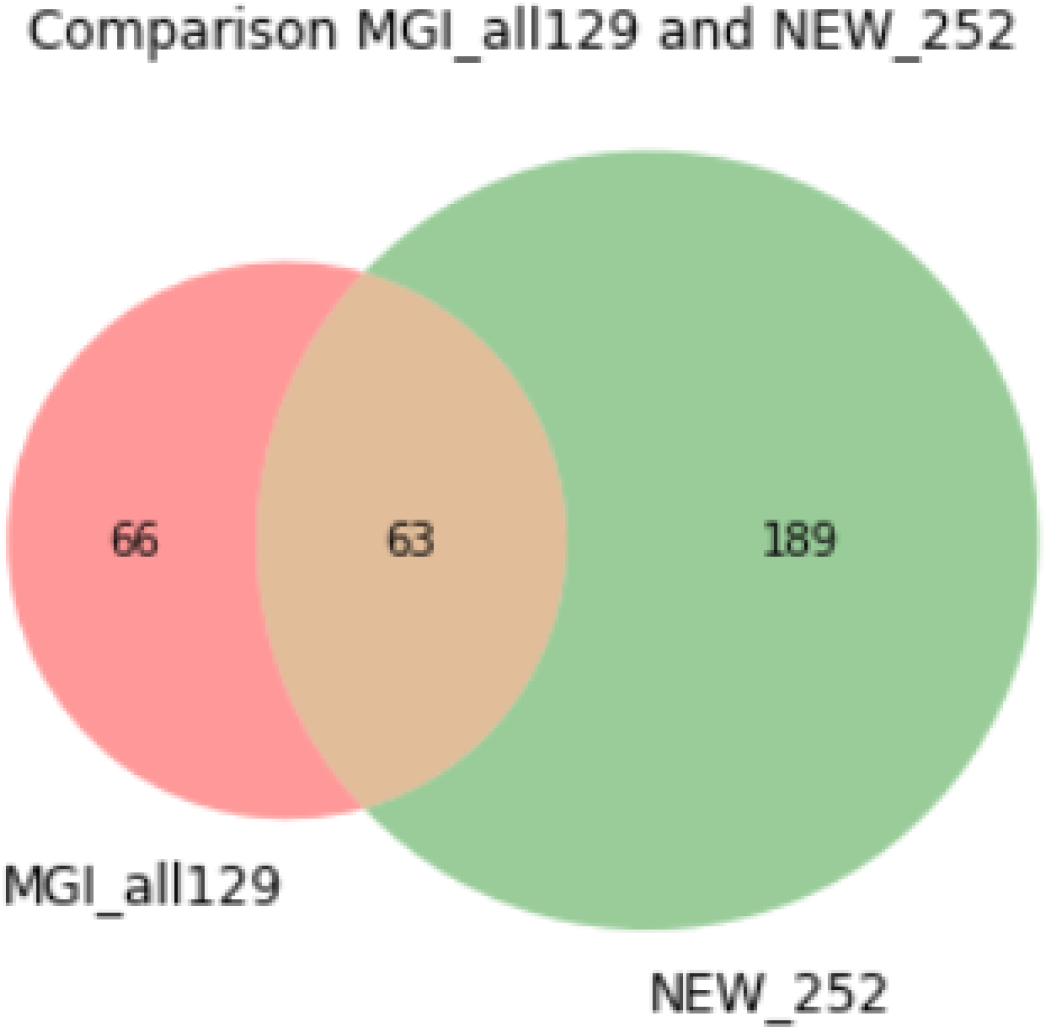
Venn diagram for comparing the sequences of custom and original MGI barcodes from the 128 barcode set and 999 validation barcode provided by MGI. Custom barcodes are shown in green, original MGI barcodes which sequences do not overlap with quads are in red, the MGI barcodes overlapping with quads (used as roots for the quads) are in orange.

**Figure 5.**
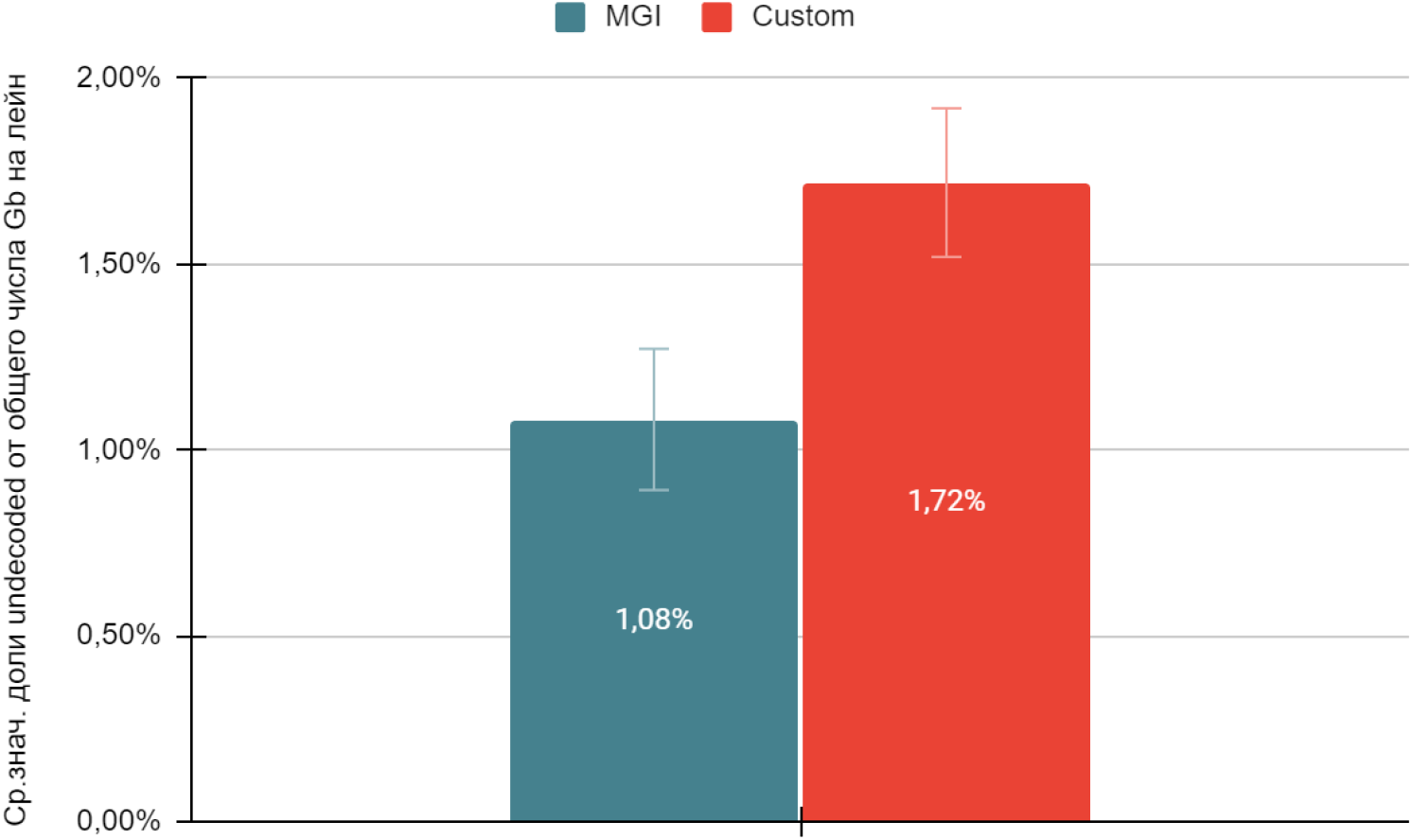
The average ratio of undecoded reads (%) and complete data (Gb) per lane for the libraries with MGI barcodes (in blue, data from 40 lanes) and with custom barcodes (red, data from 7 lanes).

Thus, we developed a viable approach for designing custom barcodes that allows for simultaneous sequencing of >96 samples on MGI called a ‘quad method’. We obtained 189 custom barcodes that can be combined with the 63 MGI barcodes to generate 63 balanced quads. One barcode from each quad is an original MGI barcode (nA, n is a number of an original barcode), the other three are custom barcodes and complement it (nB, nC, nD).

These quads can be combined with each other at any ratio and number if the ratio between the barcodes from the same quad remain equal. It is possible to create the library pools with 4n+2 barcodes, where n is a number of quads, which can include any two barcodes from the other quad. In this case, the fraction of two last barcodes should not exceed the fractions of the others. This method was validated in practice, the percentages of undecoded reads from total data in GB for lanes loaded with the libraries containing custom adapters was 1.6 times higher than in case of MGI adapters.

## Discussion

The MGI platform is designed for the fast high-throughput mass sequencing offering undeniable benefits, yet prone to limitations. We attempted to overcome certain limits resulting from the solutions and kits provided by the manufacturer. The custom barcodes we devised enable altering the ratio and the number of libraries loaded to a lane depending on the purpose and required data amount. Using the BC-store software we had earlier developed, the libraries can be more easily and quickly pooled for sequencing on the MGI machines both in the paired-end or single-end modes. Our approach allows for improving the efficiency of sequencing and expanding the possibilities of the MGI platform. However, it is important to bear in mind that the combinations of the quads with some original MGI adapters not included in the quads can fail to meet the compatibility criterion for the mismatch number. That is why we recommend to check whether they are balanced using the BC-Store software. Moreover, we assume that the percentage of useful data may be reduced in case of insufficient purity of the synthesized oligonucleotides and the prepared adapters. Therefore, taking into account all advantages and disadvantages of this method, it can be used as a complementary or alternative solution for the solution provided by MGI.

## Acknowledgements

Not applicable.

