## Supplementary material for "DESIGNING OF CUSTOM BARCODES FOR SEQUENCING ON THE MGI PLATFORM": S1 Figure

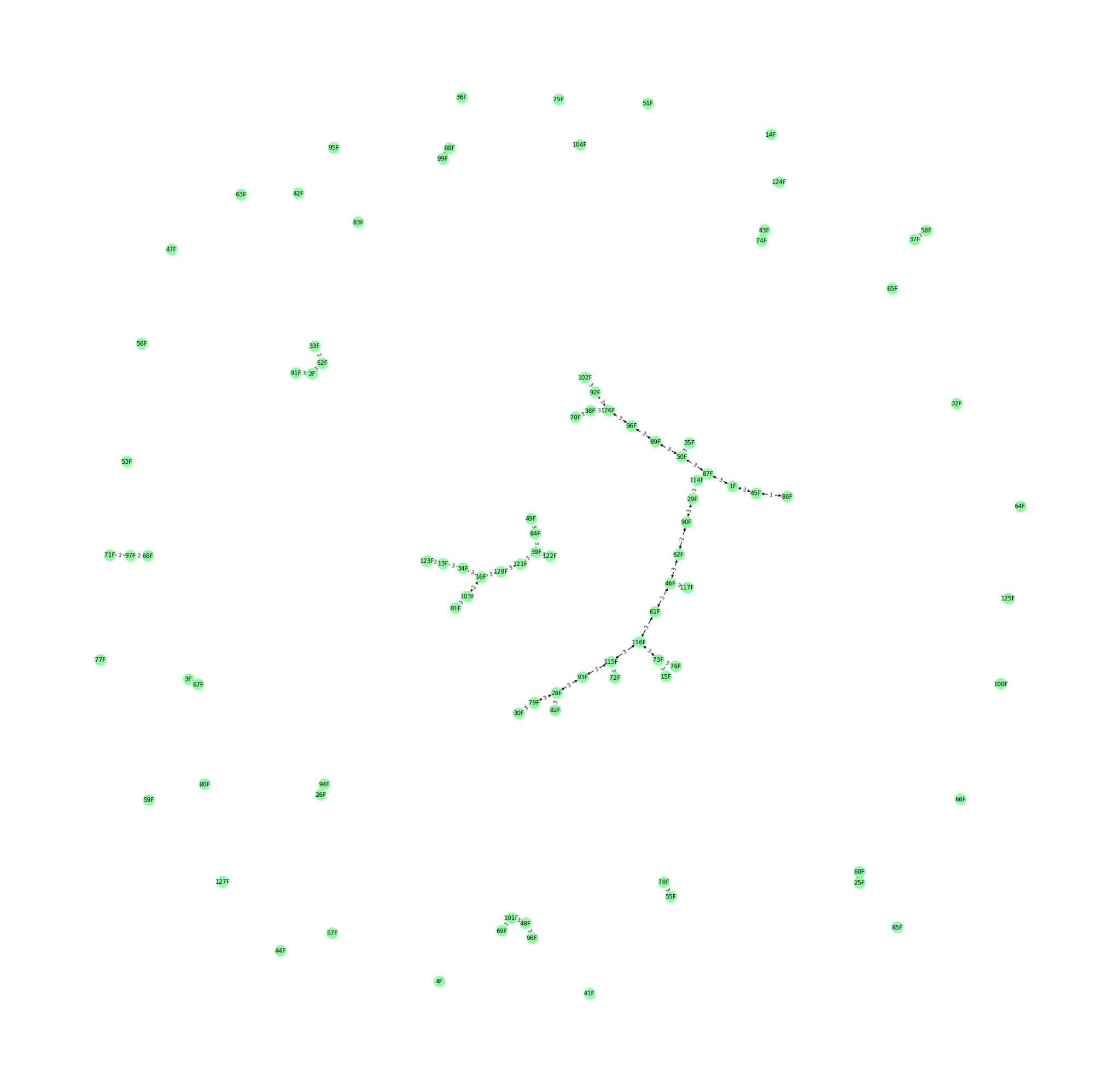

S1 Figure. A graph of incompatible quads. The incompatibility is caused by a conflict between the original MGI barcodes and generated 96 quads. An edge connects combinations with less than 4 mismatches.
