## Supplementary material for "DESIGNING OF CUSTOM BARCODES FOR SEQUENCING ON THE MGI PLATFORM": S2 Figure

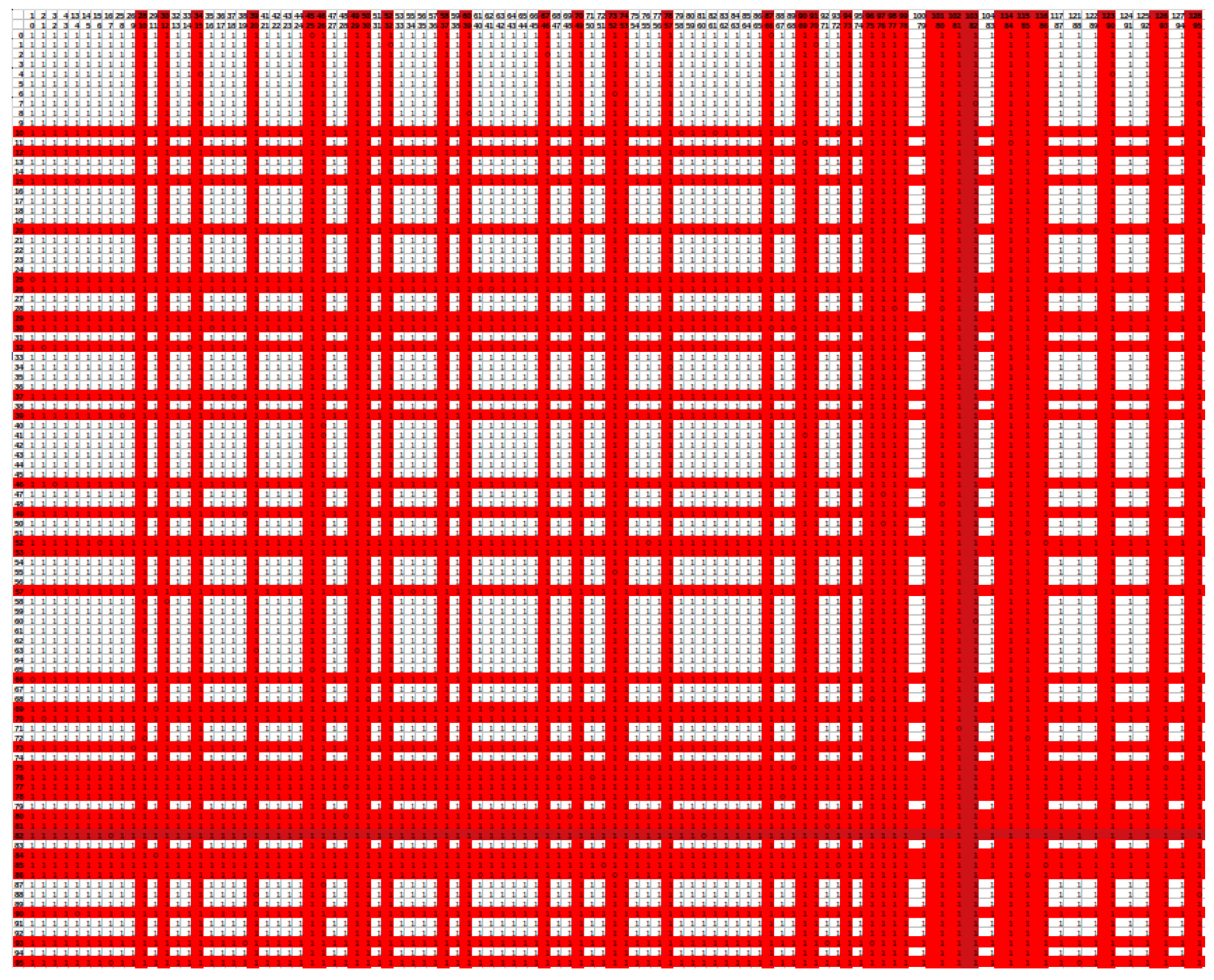

S2 Figure. An adjacency matrix based on the graph of incompatible quads. Incompatible quads are shown as 0 (contain barcodes with less than 4 mismatches), compatible quads as 1(all pairs with mismatches ≥4).
